## Supplementary Material for "Structural basis for adhesion G protein-coupled receptor Gpr126 function"

Supplementary Figures 1-7

Supplementary Tables 1-3

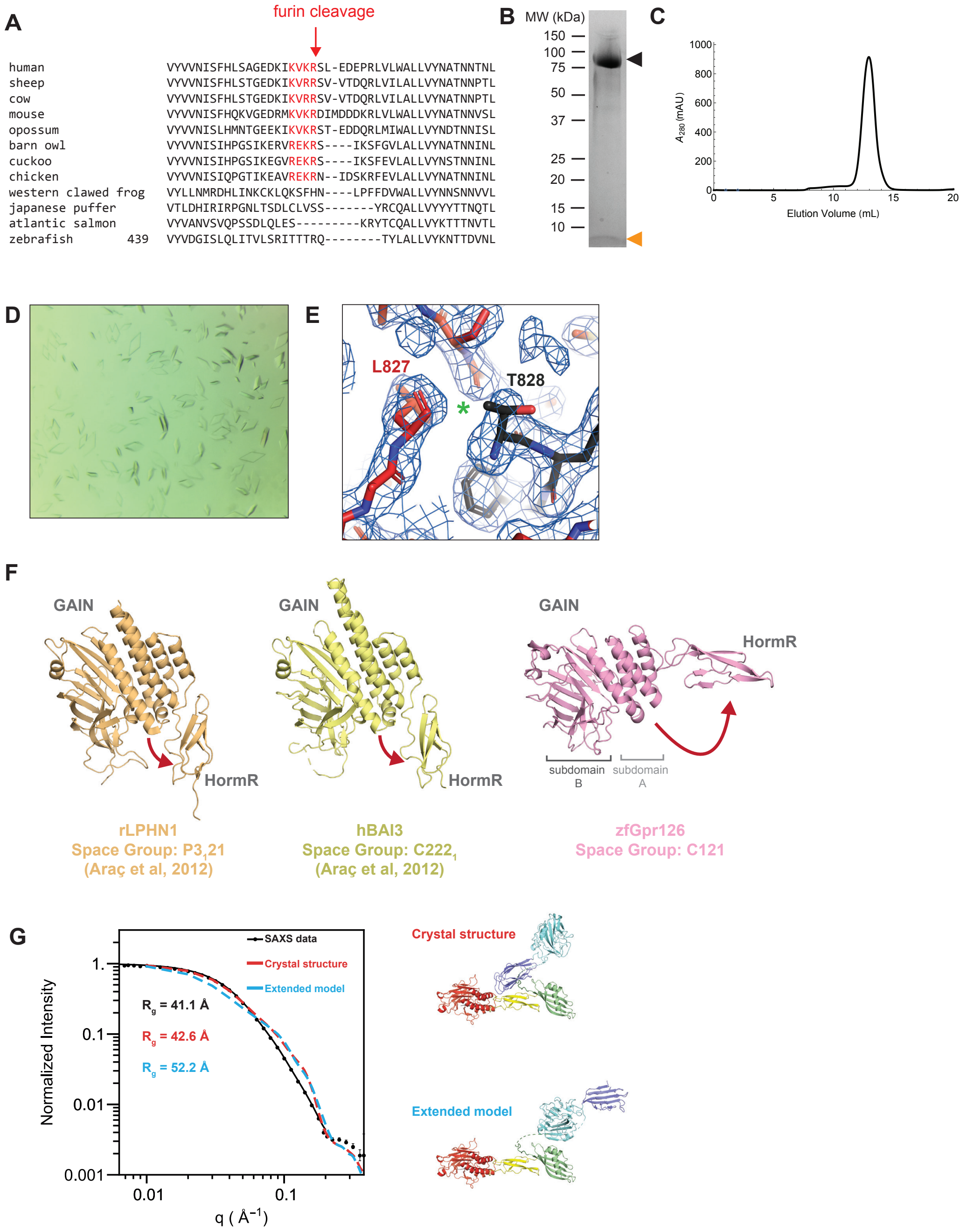


**Leon et al., Supplementary Figure 1**

**Supplementary Figure 1. Purification and crystallization of Gpr126 ECR.** (A) Sequence alignment of Gpr126 furin-cleavage site from various species. Consensus cleavage site residues are colored red. (B) SDS-PAGE of purified Gpr126 ECR showing bands for N-terminal fragment (black arrowhead) and tethered peptide (orange arrowhead). Source data are provided as a Source Data file. (C) Size exclusion column profile of purified Gpr126 ECR. (D) Crystals of Gpr126 ECR. (E) 2Fo-Fc electron density map for Gpr126 ECR showing lack of electron density at autoproteolysis site (asterisk) generating an N-terminal fragment (red) and C-terminal fragment (black). (F) HormR+GAIN domain structures and space groups of rLPHN1 (orange), hBAI3 (yellow), and zfGpr126 (pink). Red arrows indicate angles between GAIN and HormR domains, highlighting difference seen in zfGpr126 compared to all other structures. (G) Experimental SAXS curve for Gpr126 ECR (black) fit to a smooth regularized scattering curve (i.e. the Fourier transform of p(r), solid black line, see methods), simulated SAXS curve for Gpr126 ECR based on crystal structure (red), and simulated SAXS curve for extended model of Gpr126 ECR (blue). Images of crystal structure and extended model used in the analysis are shown.

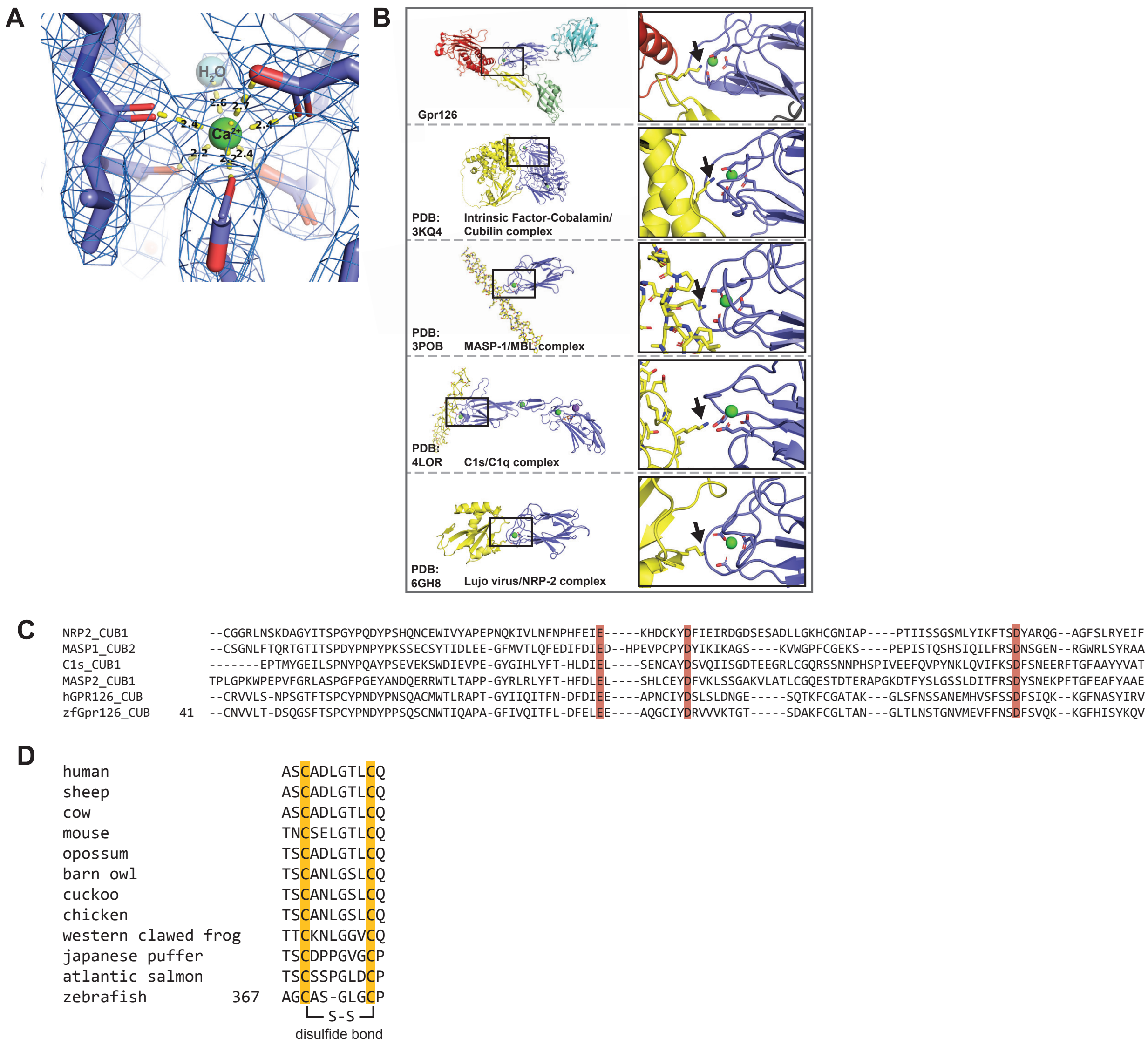


### Leon et al., Supplementary Figure 2

**Supplementary Figure 2. Analysis of the Gpr126 CUB domain.** (A) 2Fo-Fc electron density map for Gpr126 ECR showing density for calcium ion (green sphere). Coordination interactions are shown as yellow dashed lines and distances (Å) are noted. (B) (Left panels) Structures of proteins that use calcium-binding CUB domains (blue) to bind ligands (yellow). (Right panels) Close-up views of the interfaces with arrows pointing to the lysine residues on the ligands which interact with calcium-binding residues on the CUB domains. (C) Sequence alignment of CUB domains from various proteins, highlighting conserved calcium-coordinating residues (red). (D) Alignment of the disulfide-bond loop region from various species showing the cysteines (bright orange) are conserved.

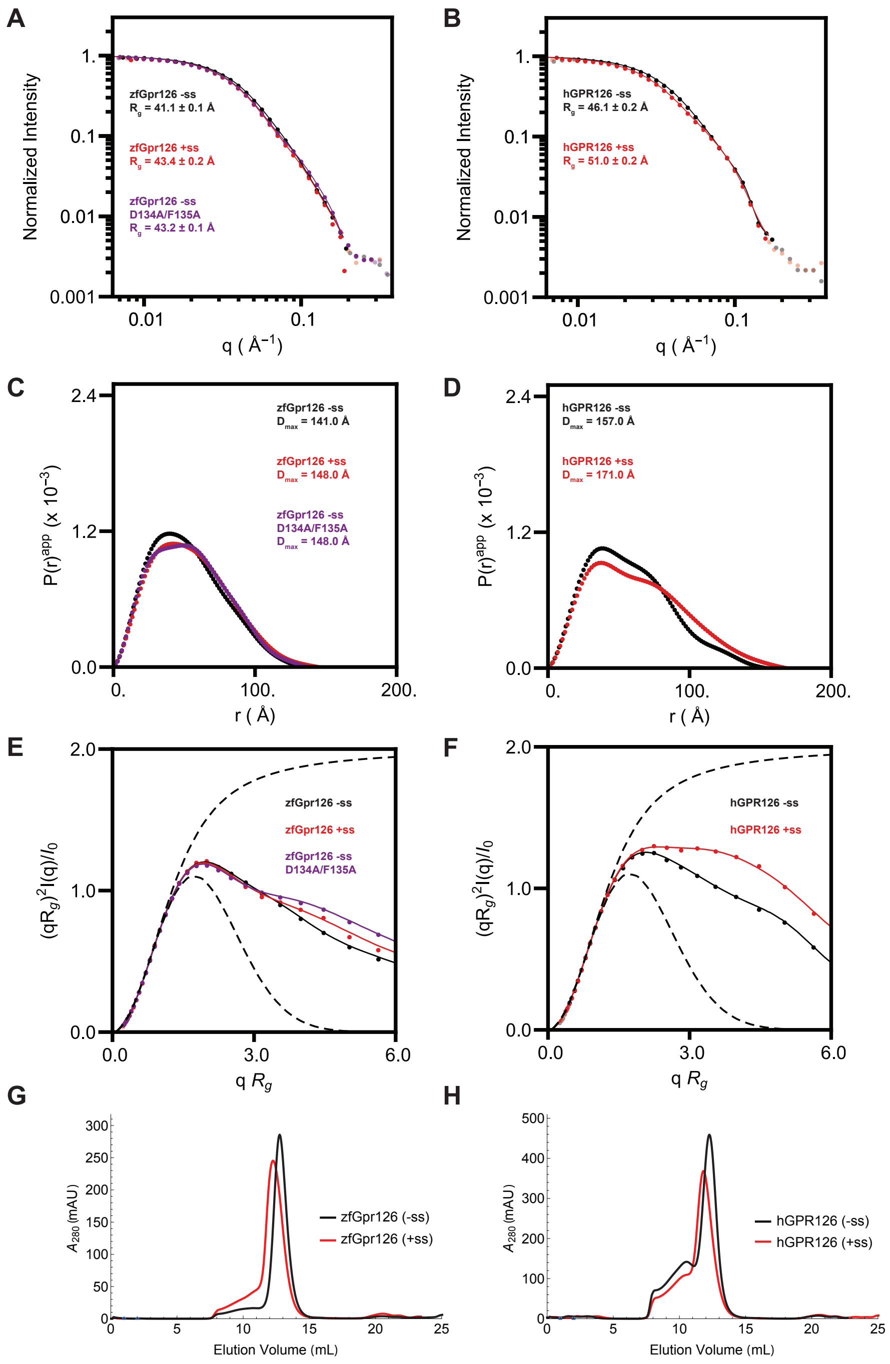


**Leon et al., Supplementary Figure 3**

**Supplementary Figure 3. Differences in ECR conformation between Gpr126/GPR126 splice isoforms.** (A) SAXS curves for zebrafish Gpr126 (-ss) (black), (+ss) (red), and (-ss) D134A/F135A (purple). (B) SAXS curves for human R468A (furin-resistant) GPR126 (-ss) (black) and (+ss) (red). (C, D) P(r) plots for zebrafish Gpr126 (C) and human R468A GPR126 (D) constructs. $P(r)^{app}$ indicates a mixed protein/carbohydrate system (glycans present). (E, F) Normalized Kratky plots for zebrafish Gpr126 (E) and human R468A GPR126 (F) constructs. Dashed lines show profiles expected for a random walk and compact (Guinier) particle located at the top and bottom, respectively of the plots. Solid lines show experimental fit with a smooth regularized scattering curve (i.e. the Fourier transform of p(r), solid black line, see methods). (G, H) Size exclusion column profiles for zebrafish Gpr126 and human R468A GPR126 splice isoforms.

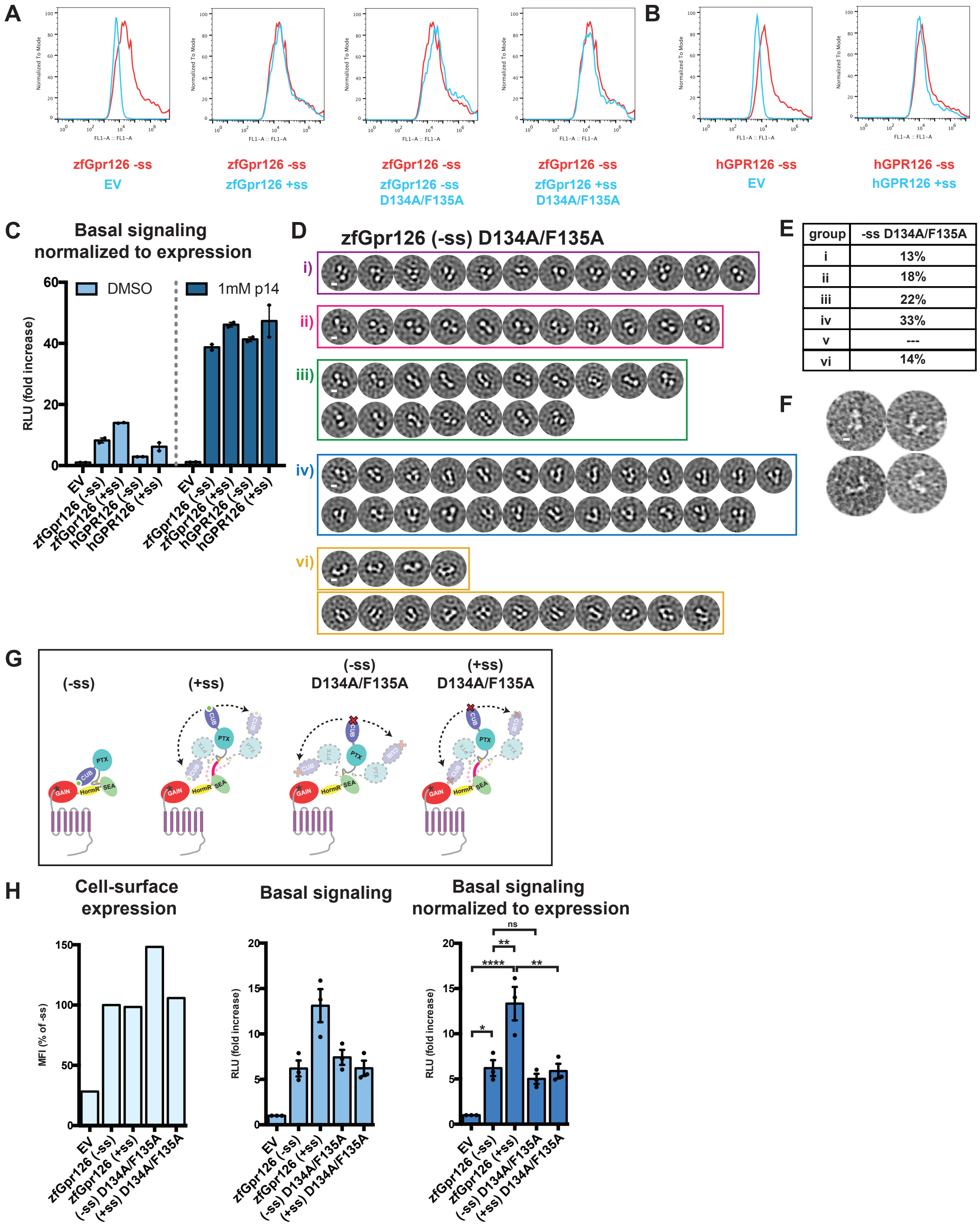


**Leon et al., Supplementary Figure 4**

**Supplementary Figure 4. Analysis of Gpr126 cAMP signaling and ECR conformation.** (A, B) Surface expression of zebrafish Gpr126 (A) and human GPR126 (B) constructs presented as overlay of typical flow cytometry histograms. HEK293 cells transfected with N-terminal FLAG-tagged full-length Gpr126/GPR126 constructs or empty vector (EV) were stained with anti-FLAG antibody without permeabilization. Expression (i.e. FLAG signal) for cells transfected with each construct (blue) is normalized to expression for cells transfected with the zebrafish or human -ss isoform (red). (C) Basal (left) and activated (right) cAMP signaling normalized to cell-surface expression for empty vector (EV), zebrafish Gpr126 splice isoforms, and human GPR126 splice isoforms. Data presented as mean ± SEM, n = 2. (D) Negative stain EM 2D class averages for zebrafish Gpr126 -ss D134A/F135A ECR constructs. Class averages are categorized according to similar orientations as in Figure 3. (i, ii, iii, iv) are observed in both -ss and +ss isoforms as well as the -ss D134A/F135A mutant. (vi) represents open-like conformations. Scale bars (white) represents 50 Å. (E) Quantification of percentage of particles per category. (F) Representative individual particles for -ss D134A/F135A. (G) ECR conformations are depicted as cartoons. The red cross represents the D134A/F135A mutation. Black arrows with dashed lines indicate dynamic ECR conformation. (H) (left) Cell-surface expression levels for EV, zebrafish Gpr126 splice isoforms, and mutant D134A/F135A isoforms, measured using flow cytometry to detect binding of anti-FLAG antibody to cells expressing FLAG-tagged Gpr126. The Gpr126 cell-surface expression levels are normalized to the control EV signal. Error bars are not shown because expression levels are presented as median fluorescence intensities of 10,000 cells for each population of transfected cells, for a single flow cytometry experiment representative of at least three independent experiments. (center) Basal signaling measured by the cAMP signaling assay. Data are shown as fold increase over EV of RLU (relative luminescence units). (right) Basal cAMP signaling normalized to cell-surface expression. ns, $P > 0.05$; *, $P \leq 0.05$; **, $P \leq 0.01$; ***, $P \leq 0.001$; ****, $P \leq 0.0001$; by one-way ANOVA and Tukey's multiple comparisons test. Signaling data are presented as mean ± SEM, n = 3, and are representative of at least three independent experiments. Source data are provided as a Source Data file.

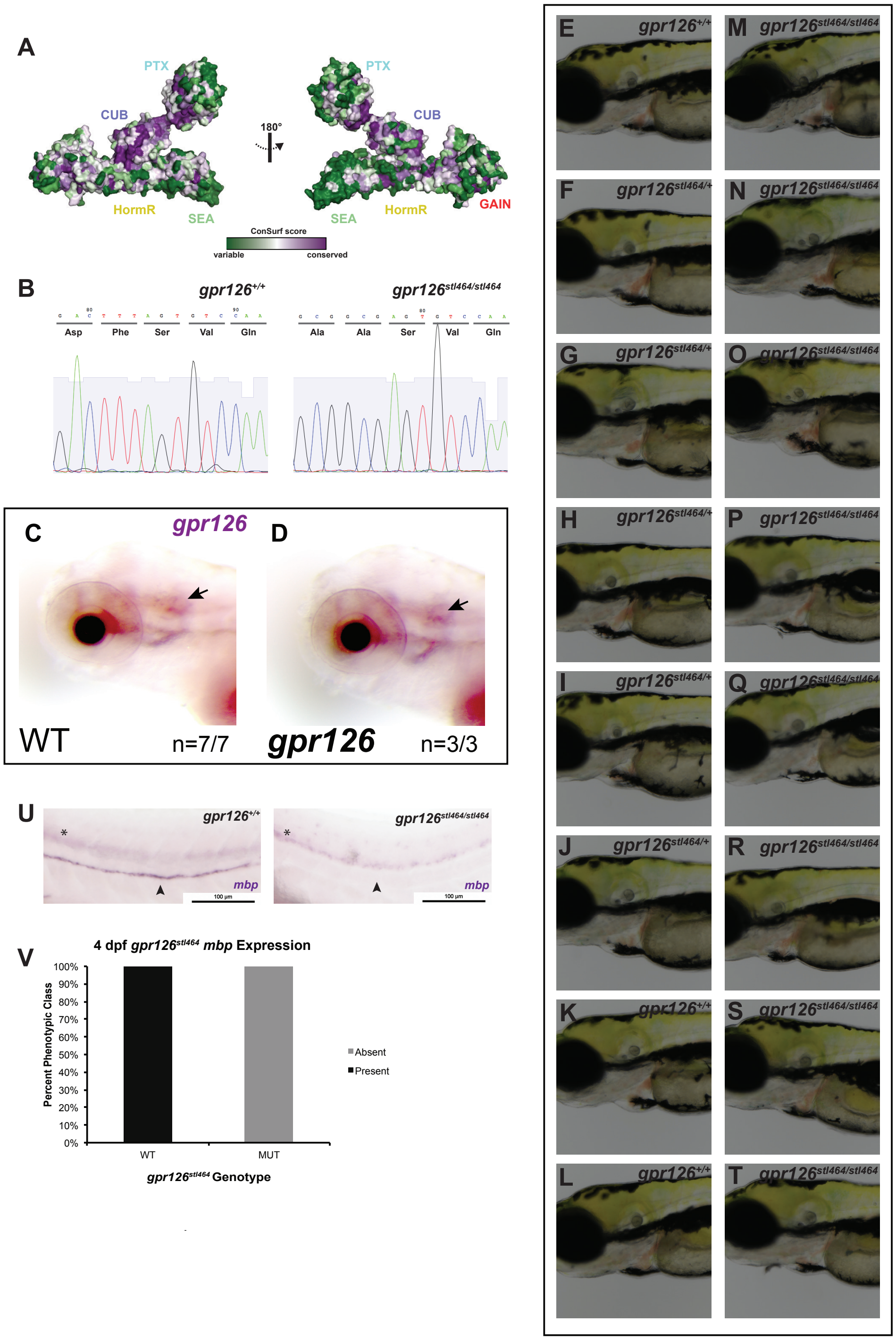


**Leon et al., Supplementary Figure 5**

**Supplementary Figure 5. Mutation of a conserved site in the ECR of Gpr126 is abolishes myelination.** (A) Conservation score of each residue mapped onto the surface of Gpr126 ECR. (B) Sequence traces for *gpr126*$^{+/+}$ *and gpr126*$^{stl464/stl464}$ zebrafish, showing D134A/F135A mutations in *gpr126*$^{stl464/stl464}$. (C, D). *In situ* hybridization shows similar expression of *gpr126* in 5 dpf wild-type siblings (C) and *gpr126*$^{stl464}$ mutants (D). Arrows denote ears. (E-L) 4 dpf wild-type and *gpr126*$^{464/+}$ larvae. (M-T) 4 dpf *gpr126*$^{464/464}$ larvae. No substantial heart defects were observed in mutants at this resolution. (U) 4 dpf wild-type larvae express *mbp* throughout the posterior lateral line nerve (PLLn, arrowhead), whereas 4 dpf *gpr126*$^{stl464/stl464}$ larva lack *mbp* expression along the PLLn (arrowhead). Asterisks indicate CNS. (V) Quantification of presence or absence of *mbp* expression in wild-type and mutant larvae.

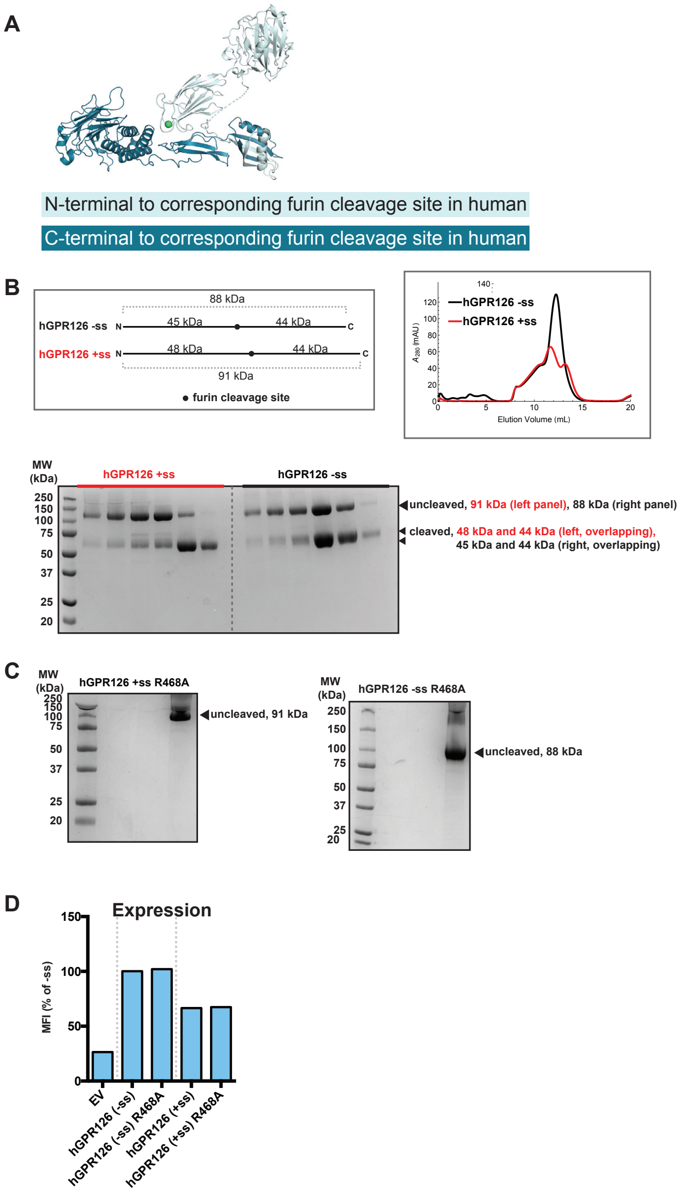


**Leon et al., Supplementary Figure 6**

**Supplementary Figure 6. Furin cleavage of GPR126.** (A) Structure of Gpr126 ECR. The region N-terminal to the corresponding human furin cleavage site is colored light blue and the region C-terminal to the corresponding human furin cleavage site is colored dark blue. (B) (top, left) Molecular weights for full-length ECR and furin-cleaved ECR fragments of human GPR126 -ss and +ss isoforms are shown, with the furin cleavage site shown as a black circle. (top, right) Size exclusion column profiles for human GPR126 -ss (black) and +ss (red) ECR proteins. (bottom) SDS-PAGE analysis of size exclusion column fractions, showing partial cleavage by furin of both isoforms. The top bands (~88, 91 kDa) correspond to uncleaved GPR126 -ss and +ss ECR, respectively. The bottom bands (~45/44 and 48/44 kDa) correspond to cleaved GPR126 -ss and +ss ECR, respectively. The N-terminal and C-terminal cleaved fragments are close in size and overlap on the gel. (C) SDS-PAGE analysis of human GPR126 -ss (left) and +ss (right) ECR constructs with furin-resistant R468A mutations, showing loss of cleaved fragments. Source data are provided as a Source Data file. (D) GPR126 cell-surface expression levels on HEK293 cells transfected with empty vector (EV) and various human GPR126 constructs. Error bars are not shown because expression levels are presented as median fluorescence intensities of 10,000 cells for each population of transfected cells, for a single flow cytometry experiment representative of at least three independent experiments.

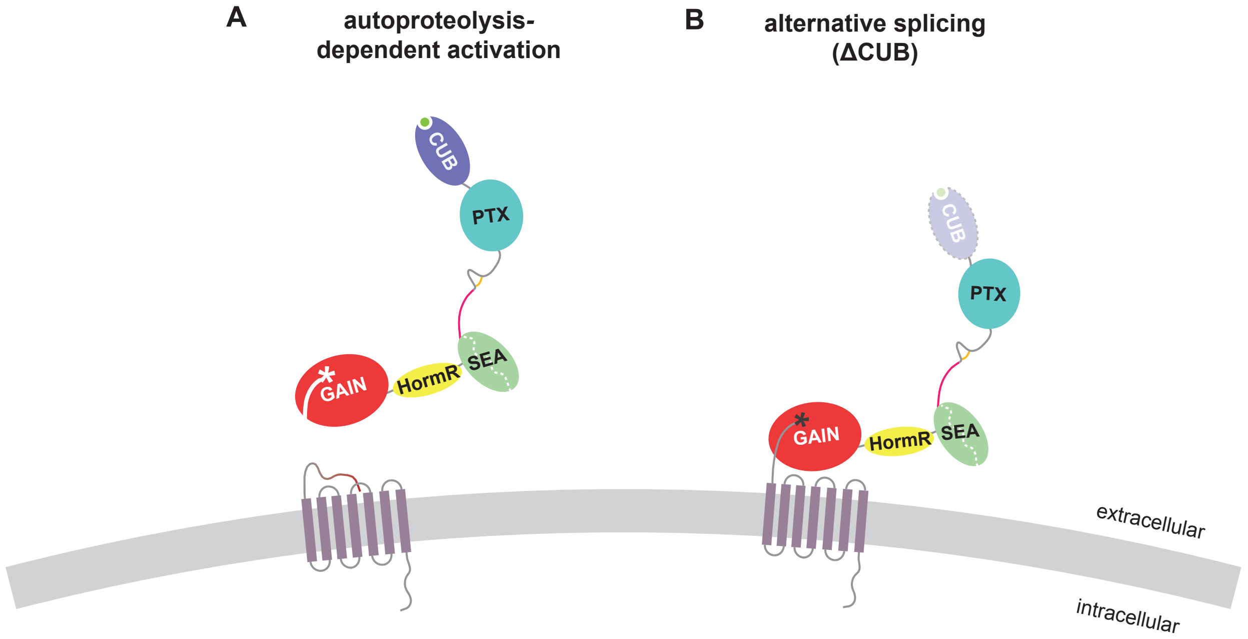


**Leon et al., Supplementary Figure 7**

**Supplementary Figure 7. Additional models for activation and regulation of Gpr126 function.** (A) Mechanical force exerted on the Gpr126 ECR leads to exposure of tethered peptide and activation of the receptor. (B) An alternative splice isoform lacks the CUB domain and may be a mechanism for receptor regulation.

**Supplementary Table 1. Data Collection and Refinement Statistics**

| Protein | Gpr126 ECR Native Dataset | Gpr126 ECR SeMet SAD |
|---|---|---|
| Data Collection | | |
| Integration Package | HKL2000 | HKL2000 |
| Wavelength | 1.033 Å | 0.979 Å |
| Space group | C2 | C2 |
| Cell dimensions | | |
| *a*, *b*, *c* | 144.96 Å, 59.45 Å, 168.39 Å | 145.20 Å, 59.33 Å, 168.72 Å |
| α, β, γ | 90°, 107.82°, 90° | 90°, 108.16°, 90° |
| Resolution | 50–2.40 Å (2.46–2.40 Å) | 50–2.34 Å (2.38–2.34 Å) |
| $R_{sym}$ or $R_{merge}$ | 0.064 (0.664) | 0.086 (0.492) |
| $CC_{1/2}$, highest res. bin | 0.557 | 0.376 |
| $I/\sigma I$ | 15.0 (0.9) | 13.0 (1.4) |
| Completeness | 77.9% (25.4%) | 71.6% (9.7%) |
| Redundancy | 2.9 (1.7) | 3.8 (1.3) |
| Number of measured reflections | 125687 | 157602 |
| Number of unique reflections | 43044 | 41385 |
| Refinement Statistics | | |
| $R_{work}/R_{free}$ | 0.216/0.271 | |
| Number of atoms | | |
| Protein | 5844 | |
| Water | 146 | |
| Other | 169 | |
| Average B-factors | | |
| Protein | 28.8 $Å^2$ | |
| Water | 26.1 $Å^2$ | |
| Other | 57.5 $Å^2$ | |
| Rmsds | | |
| Bond lengths | 0.009 Å | |
| Bond angles | 1.118° | |
| Ramachandran plot statistics | | |
| Most favorable | 90.98% | |
| Allowed | 8.22% | |
| Disallowed | 0.8% | |

**Supplementary Table 2. Information for various Gpr126 species**

| class | species | common name | furin cleavage site | evidence of splice site between PTX/SEA | two cysteines present in loop |
|---|---|---|---|---|---|
| mammalia | Ailuropoda_melanoleuca | gianta panda | yes | yes | yes |
| reptilia | Alligator_mississippiensi | American alligator | yes | yes | yes |
| aves | Anas_platyrhynchos | Mallard | yes | yes | yes |
| reptilia | Anolis carolinensis | green anole [lizard] | yes | yes | yes |
| aves | Antrostomus_carolinensis | Chuck-wills-widow | yes | no | yes |
| actinopterygii | Aphyosemion striatum | Red-striped Killifish | no | no | no |
| aves | Aptenodytes_forsteri | Emperor penguin | yes | no | yes |
| actinopterygii | astyanax mexicanus | mexican tetra | no | yes | yes |
| actinopterygii | Austrofundulus limnaeus | annual killifish | no | yes | no |
| mammalia | Bos_mutus | Domestic yak | yes | yes | yes |
| mammalia | Bos_taurus | cow | yes | yes | yes |
| aves | Buceros_rhinoceros_silvestris | Rhinoceros hornbill | no | no | yes |
| mammalia | Callithrix_jacchus | Marmoset | yes | yes | yes |
| aves | calypte anna | anna's hummingbird | yes | no | yes |
| mammalia | Camelus ferus | wild bactrian camel | yes | yes | yes |
| mammalia | Canis_lupus_familiaris | dog | yes | yes | yes |
| aves | Charadrius_vociferus | Killdeer | yes | no | yes |
| aves | Chlamydotis_macqueenii | Macqueen’s bustard | yes | no | yes |
| mammalia | Chlorocebus_sabaeus | green monkey | yes | yes | yes |
| aves | Colius_striatus | Speckled mouse-bird | yes | no | yes |
| aves | Corvus_brachyrhynchos | American crow | yes | yes | yes |
| mammalia | cricetulus griseus | chinese hamster | yes | yes | yes |
| reptilia | Crocodylus porosus | Australian saltwater crocodile | yes | yes | yes |
| aves | Cuculus_canorus | cuckoo | yes | no | yes |
| actinopterygii | danio_rerio | zebrafish | no | yes | yes |
| mammalia | Delphinapterus leucas | beluga whale | yes | yes | yes |
| mammalia | Dipodomys_ordii | Ord’s kangaroo rat | yes | yes | yes |
| aves | Egretta_garzetta | little egret | n/a | no | yes |
| mammalia | Equus_caballus | Horse | yes | yes | yes |

| | | | | | |
|---|---|---|---|---|---|
| mammalia | Erinaceus_europaeus | european hedgehog | yes | yes | yes |
| aves | Ficedula_albicollis | Collared flycatcher | yes | yes | yes |
| actinopterygii | Fundulus_heteroclitus | Mummichog | no | yes | yes |
| aves | Gallus_gallus | chicken | yes | yes | yes |
| mammalia | Gorilla_gorilla_gorilla | gorilla | yes | yes | yes |
| aves | Haliaeetus_albicilla | White-tailed eagle | yes | no | yes |
| mammalia | Heterocephalus_glaber | Naked mole-rat | yes | yes | yes |
| mammalia | homo_sapiens | human | yes | yes | yes |
| actinopterygii | Ictalurus punctatus | channel catfish | no | yes | yes |
| actinopterygii | Larimichthys_crocea | Large yellow croaker | no | yes | yes |
| actinopterygii | Lates calcarifer | baramundi perch | no | yes | yes |
| actinopterygii | Lepisosteus_oculatus | Spotted gar | yes | yes | yes |
| mammalia | Loxodonta_africana | African bush elephant | yes | yes | yes |
| mammalia | Macaca fascicularis | crab-eating macaque | yes | yes | yes |
| mammalia | Macaca_mulatta | Rhesus macaque | yes | yes | yes |
| aves | Manacus_vitellinus | Golden-collared manakin | no | no | yes |
| aves | Meleagris_gallopavo | Turkey | yes | no | yes |
| aves | Merops nubicus | northen carmine bee-eater | n/a | no | yes |
| mammalia | Mesocricetus auratus | golden hamster | yes | yes | yes |
| mammalia | Monodelphis_domestica | opossum | yes | yes | yes |
| mammalia | Mus_musculus | mouse | yes | yes | yes |
| mammalia | Mustela_putorius_furo | Ferret | yes | yes | yes |
| mammalia | Myotis_davidii | Vesper bat | yes | no | yes |
| mammalia | Myotis_lucifugus | Little brown bat | yes | no | yes |
| aves | Nipponia_nippon | Crested ibis | no | no | yes |
| mammalia | Nomascus_leucogenys | Northern white-cheeked gibbon | yes | yes | yes |
| actinopterygii | Nothobranchius kuhntae | Beira killifish | no | no | no |
| actinopterygii | Nothobranchius_furzeri | Turquoise killifish | no | yes | no |
| actinopterygii | Oncorhynchus kisutch | coho salmon | no | yes | yes |
| actinopterygii | Oncorhynchus mykiss | rainbow trout | no | yes | yes |
| actinopterygii | Oncorhynchus tshawytscha | chinook salmon | no | yes | yes |
| aves | Opisthocomus_hoazin | hoatzin bird | yes | no | yes |
| actinopterygii | Oreochromis_niloticus | Nile tilapia | no | yes | yes |
| mammalia | Ornithorhynchus_anatinus | Platypus | yes | no | yes |

| | | | | | |
|---|---|---|---|---|---|
| mammalia | Oryctolagus_cuniculus | European rabbit | yes | yes | yes |
| mammalia | Otolemur_garnettii | Northern great galago | yes | yes | yes |
| mammalia | Ovis_aries | sheep | yes | yes | yes |
| mammalia | Pan_troglodytes | chimpanzee | yes | yes | yes |
| mammalia | Panthera pardus | leopard | yes | yes | yes |
| mammalia | Papio_anubis | Olive baboon | yes | yes | yes |
| aves | Pelecanus_crispus | Dalmatian pelican | yes | no | yes |
| reptilia | Pelodiscus_sinensis | Chinese soft-shell turtle | no | yes | yes |
| aves | Phaethon_lepturus | White-tailed tropicbird | yes | no | yes |
| aves | Phalacrocorax_carbo | great cormorant | no | no | yes |
| aves | Picoides_pubescens | downy woodpecker | no | no | yes |
| aves | Podiceps_cristatus | great crested grebe | yes | no | yes |
| actinopterygii | Poecilia_formosa | Amazon Molly | no | yes | yes |
| actinopterygii | Poeciliopsis prolifica | blackstripe livebearer | no | yes | yes |
| mammalia | Pongo_abelii | Sumatran orangutan | yes | yes | yes |
| aves | Pterocles_gutturalis | Yellow-throated sandgrouse | yes | no | yes |
| mammalia | Pteropus_alecto | Black flying fox | yes | no | yes |
| aves | Pygoscelis_adeliae | Adelie penguin | no | no | yes |
| mammalia | Rattus_norvegicus | Brown rat | yes | yes | yes |
| actinopterygii | Salmo_salar | atlantic salmon | no | yes | yes |
| mammalia | Sarcophilus_harrisii | tasmanian devil | no | yes | yes |
| mammalia | sus scrofa | pig | yes | yes | yes |
| aves | Taeniopygia_guttata | Zebra finch | yes | no | yes |
| actinopterygii | Takifugu_rubripes | japanese puffer | no | yes | yes |
| aves | Tauraco_erythrolophus | Red-crested turaco | no | no | yes |
| reptilia | Terrapene mexicana triunguis | Three-toed box turtle | yes | yes | yes |
| actinopterygii | Tetraodon_nigroviridis | green pufferfish | no | yes | yes |
| aves | Tinamus_guttatus | White-throated tinamou | no | no | yes |
| aves | Tyto_alba | barn owl | yes | no | yes |
| amphibia | Xenopus_tropicalis | western clawed frog | n/a | yes | yes |
| actinopterygii | Xiphophorus_maculatus | southern platyfish | n/a | yes | yes |

**Supplementary Table 3. SAXS analysis for Gpr126/GPR126 constructs**

(*a*) Sample details

| | zfGpr126 -ss ECR | zfGpr126 +ss ECR | zfGpr126 -ss ECR D134A/F135A | hGPR126 -ss ECR R468A | hGPR126 +ss ECR R468A |
|---|---|---|---|---|---|
| Organism | Zebrafish | Zebrafish | Zebrafish | Human | Human |
| Source (Catalogue No. or reference) | Monk Lab | Monk Lab | Monk Lab | Addgene plasmid #66314 | Addgene plasmid #66314 |
| Description | C6KFA3 splice variant S2, C-terminal 8xHis tag | C6KFA3 splice variant S1, C-terminal 8xHis tag | C6KFA3 splice variant S2 with D134A and F135A mutations, C-terminal 8xHis tag | Q86SQ4, C-terminal 8xHis tag, R468A mutation | Q86SQ4, C-terminal 8xHis tag, R468A mutation |
| Extinction coefficient ε (A280, 0.1%(w/v)) | 1.094 | 1.066 | 1.096 | 1.195 | 1.155 |
| Partial specific volume $\bar{\bar{v}}$ ($cm^3\ g^{-1}$) | 0.733 | 0.734 | 0.733 | 0.735 | 0.734 |
| Mean solute and solvent scattering length densities and mean scattering contrast $\Delta\bar{\bar{\rho}}$ ($10^{10}\ cm^{-2}$) | 2.892 (12.352 – 9.460) | 2.886 (12.346 – 9.460) | 2.891 (12.351 – 9.460) | 2.874 (12.334 – 9.460) | 2.881 (12.341 – 9.460) |
| Molecular mass *M* from chemical composition (Da) | 86371 | 88708 | 86251 | 88444 | 91467 |
| SEC-SAXS column | Superdex 200 10/300 | | | | |
| Loading concentration, (mg $ml^{-1}$) | 5.0 | 2.7 | 2.5 | 4.2 | 4.2 |
| Injection volume (µl) | 200 | 200 | 200 | 200 | 200 |
| Flow rate (ml $min^{-1}$) | 0.7 | 0.7 | 0.7 | 0.7 | 0.7 |
| Solvent composition and source | 20 mM HEPES pH 7.2, 150 mM NaCl | | | | |

(*b*) SAS data collection parameters

| | |
|---|---|
| Source, instrument and description or reference | Advanced Photon Souce at Argonne National Labs, BioCAT 18-ID with Pilatus 3 1M detector |
| Wavelength (Å) | 1.03 |
| Beam size (µm) | 172 x 172 |
| Sample-to-detector distance (m) | 1.0 |
| *q*-measurement range ($Å^{-1}$) | 0.00540153 – 0.383203 |
| Absolute scaling method | N/A |
| Basis for normalization to constant counts | Incident flux |

| | |
|---|---|
| Method for monitoring radiation damage, X-ray dose where relevant | Dose < $10^{12}$ photons $sec^{-1}$. |
| Exposure time, number of exposures | Continuous 0.8 s data-frame measurements of SEC elution |
| Sample configuration including path length and flow rate where relevant | In-line SEC-SAXS, pathlength = 1.5 mm at 0.7 ml $min^{-1}$. |
| Sample temperature (ºC) | 22 |

(*c*) Software employed for SAS data reduction, analysis and interpretation

| | |
|---|---|
| SAS data reduction to sample–solvent scattering, and extrapolation, merging, desmearing *etc.* as relevant | BioXTAS RAW[1] |
| Calculation of ε from sequence | ProtParam[2] |
| Calculation of $\bar{\bar{v}}$ and $\Delta\bar{\bar{\rho}}$ values from chemical composition | MULCh[3] |
| Basic analyses: Guinier, *P*(*r*), scattering particle volume (*e.g.* Porod volume $V_P$ or volume of correlation $V_c$) | BioXTAS RAW[1]<br>AUTORG, DATGNOM from ATSAS[4] |
| Shape/bead modelling | N/A |
| Atomic structure modelling (homology, rigid body, ensemble) | CRYSOL[5] |
| Modelling of missing sequence from PDB files | N/A |
| Molecular graphics | The PyMOL Molecular Graphics System, Version 2.0 Schrödinger, LLC |

(*c*) Structural parameters

| | zfGpr126 -ss ECR | zfGpr126 +ss ECR | zfGpr126 -ss ECR D134A/F135A | hGPR126 -ss ECR R468A | hGPR126 +ss ECR R468A |
|---|---|---|---|---|---|
| Guinier Analysis | | | | | |
| *I*(0) | 395.1964 ± 0.8711 | 90.2641 ± 0.4052 | 153.6211 ± 0.4943 | 124.1835 ± 0.4630 | 129.0501 ± 0.5463 |
| $R_g$ (Å) | 40.1790 ± 0.1624 | 42.4661 ± 0.2703 | 42.8000 ± 0.2312 | 44.0908 ± 0.2382 | 48.9428 ± 0.2815 |
| *q*-range ($Å^{-1}$) | 0.0054 – 0.02957 | 0.00642 – 0.03119 | 0.00642 – 0.02843 | 0.00642 – 0.02935 | 0.0052 – 0.02721 |
| Coefficient of correlation, $R^2$ | 0.9937 | 0.9840 | 0.9902 | 0.9704 | 0.9737 |
| *P*(*r*) analysis | | | | | |
| *I*(0) | 398.100 ± 0.6880 | 90.7600 ± 0.3192 | 153.9000 ± 0.4063 | 126.1000 ± 0.3874 | 130.7000 ± 0.4821 |
| $R_g$ (Å) | 41.130 ± 0.098 | 43.400 ± 0.189 | 43.240 ± 0.144 | 46.070 ± 0.193 | 50.980 ± 0.225 |
| $d_{max}$ (Å) | 141 | 148 | 148 | 157 | 171 |
| *q*-range ($Å^{-1}$) | 0.0076 – 0.1991 | 0.0087 – 0.1890 | 0.0052 – 0.3663 | 0.0102 – 0.1777 | 0.0076 – 0.1638 |
| Total estimate from GNOM | 0.915 | 0.825 | 0.9051 | 0.804 | 0.755 |
| Porod volume ($Å^{-3}$) | 169000 | 191000 | 181000 | 199000 | 213000 |

(*d*) Deposition IDs

| | zfGpr126 -ss ECR | zfGpr126 +ss ECR | zfGpr126 -ss ECR D134A/F135A | hGPR126 -ss ECR R468A | hGPR126 +ss ECR R468A |
|---|---|---|---|---|---|
| SASBDB ID | SASDFT9 | SASDFU9 | SASDFV9 | SASDFW9 | SASDFX9 |
